# Dual-Resonance Shapes REM Sleep: *A Mechanistic Link Between Homeostasis, Circadian Phase, and Sleep Deficit*

**DOI:** 10.1101/2025.05.02.651911

**Authors:** Irina V. Zhdanova, Vasili Kharchenko

## Abstract

How circadian and homeostatic processes interact to shape REM sleep architecture remains unresolved. Here, we empirically confirm the existence of a conserved resonance curve governing REM episode duration, as previously predicted by the wave model of sleep dynamics. This curve emerges from two converging resonance states **–** homeostatic and circadian **–** that form a stable framework across conditions. Their relative phase and amplitude determine REM structure, producing bell-shaped, linear, or bimodal profiles. The model reproduces REM dynamics during regular sleep, extension, and post-deprivation recovery. Crucially, REM duration alone provides an experimentally accessible readout: simultaneously revealing circadian phase and quantifying sleep deficit as additional hours needed to restore homeostatic equilibrium. Together, these findings reposition REM duration from a descriptive to a diagnostic marker, and a mechanistic signal, linking sleep architecture to the dynamic interplay of internal regulatory systems.

**Summary sentence:** REM episode duration emerges as a dual readout of circadian phase and sleep deficit, reflecting the interaction of homeostatic and circadian resonance processes.

---

Sleep unfolds in repeating cycles of non-REM (NREM) and REM episodes, each with distinct duration and intensity. While these patterns are well-characterized, their mechanistic origin **–** particularly for REM sleep **–** remains elusive. Among the most puzzling features is the shape of the REM episode duration curve across the sleep period, which can appear bell-shaped, linear, or even bimodal, depending on context^1-6^. This contrasts with other principal sleep measures **–** NREM episode duration, NREM intensity, and REM density **–** that retain consistent temporal profiles across sleep, emphasizing that the dynamic variability in REM episode duration is unique among these features.

Although both homeostatic sleep need and circadian timing are known to regulate REM episode duration^3^, no unified model has explained how these two systems interact to shape its dynamics.

In this study, we build on the previously introduced wave model of sleep dynamics^1^, which represents sleep and wake as interacting probability waves and captured, with high mathematical precision, key structural features of sleep under both normal and perturbed conditions^1^.

In conventional sleep science, prolonged wakefulness increases *sleep need*, typically indexed by rising sleepiness and, upon sleep onset, elevated slow-wave activity in NREM sleep^7^. A complementary concept from cognitive performance studies **–** *wake-state instability* **–** describes the increasing variability in attention and reaction time during sleep deprivation, reflecting a growing physiological difficulty in maintaining stable wakefulness^8,9^. While distinct in origin, this behavioral concept captures a similar principle to what the wave model formalizes more broadly: a rising *systemic instability* as the organism moves away from homeostatic equilibrium.

In the model (Fig. 1), this state instability maps onto the energy structure of a Morse potential **–** a formal framework that represents sleep architecture as a sequence of quantized cycles^1^. In this context, “energy” is a measure of systemic instability **–** consistent with its physical interpretation, where higher energy typically reflects a more unstable or transition-prone state. This energy increases during sustained wakefulness and declines as the system progresses through successive sleep cycles. At the model’s homeostatic equilibrium region **–** where the two potentials intersect **–** the energies of the sleep and wake states are equal, and their interaction is at its maximum.

**Figure 1.**
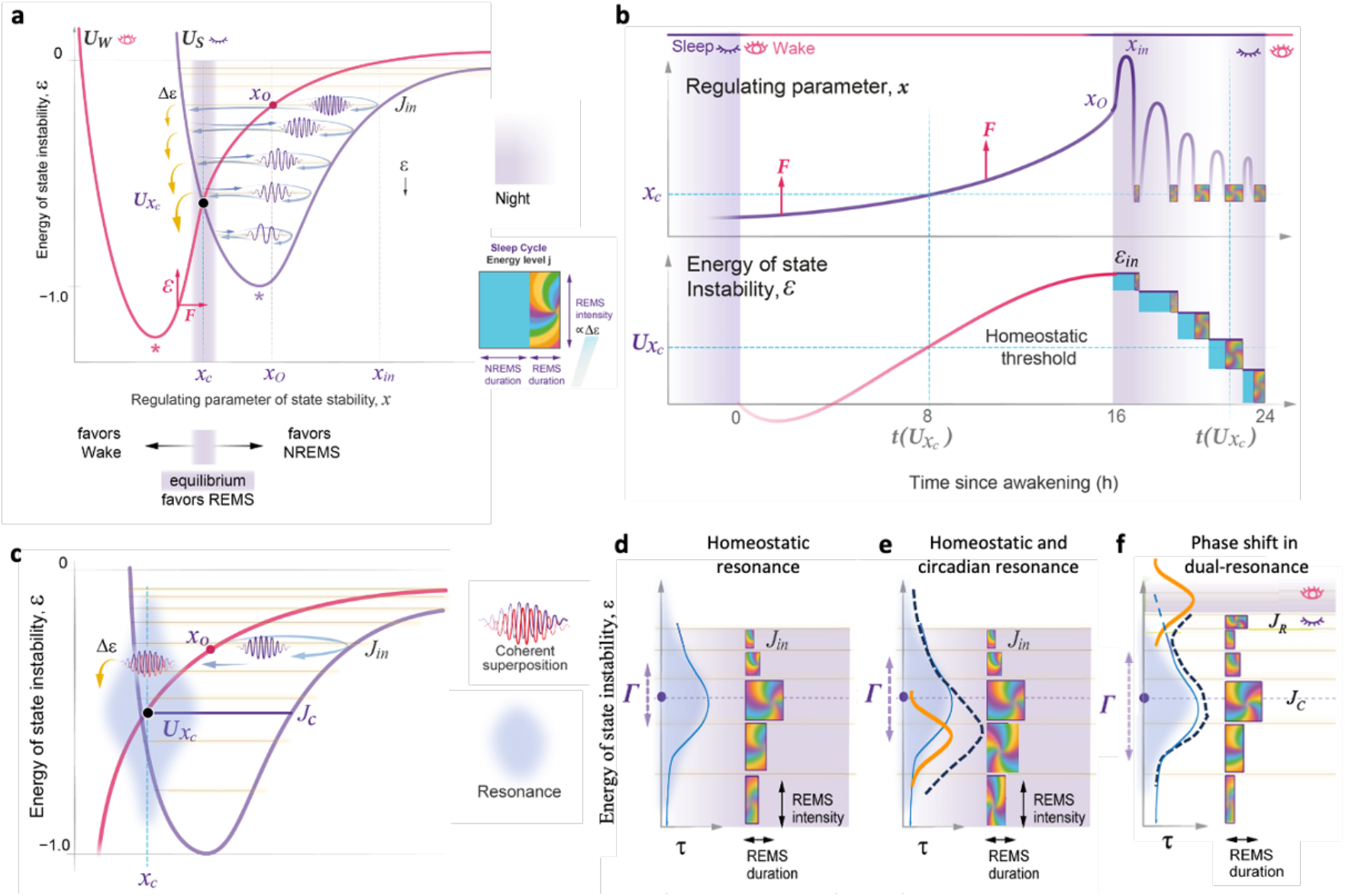
How Sleep and Wake States Interact to Shape NREM and REM Architecture. The wave model of sleep dynamics describes sleep-wake cycle as the movement of a wavepacket through a potential energy landscape^1^. **(a)** The sleep state is modeled by a Morse potential (*U*_*S*_, purple) and wake as an interacting potential (*U*_*W*,_ red). A driving force (F) during wake increases a regulatory parameter *x*, elevating state instability ε within *U*_*W*_, crossing the equilibrium threshold *x*_c_ and initiating sleep at *x*_*0*_. During sleep, the wavepacket descends through quantized energy levels and transits back to *U*_*W*_ below equilibrium (U*x*_c_). Each NREM episode’s duration and intensity correspond to the period and amplitude of a single oscillation of *x*, respectively. The energy of state instability (ε) decreases in a stepwise manner, with REM sleep intensity corresponding to the energy gaps (Δε). **(b)** Time-dependent changes in *x* and ε. Purple areas — night. **(c)** Near the crossing point of the two potentials (U*x*_c_), strong interaction gives rise to a resonance state (gray zone) at resonance level *J*_*C*_, enabling a coherent superposition of sleep and wake waves — corresponding to REM sleep. **(d)** The duration of coherent superposition (τ) and thus each REM episode duration depends on the resonance strength at that energy level, with the homeostatic resonance peak at *J*_*C*_ producing a bell-shaped REM profile across cycles. **(e)** An independent circadian resonance (orange) can further enhance the duration of REM episodes. The temporal relationship between homeostatic and circadian resonance states defines the shape and amplitude of the composite resonance state (black dashed line), as well as the dynamics of REM episode duration. **(f)** Misalignment between homeostatic and circadian resonance peaks can alter the shape of the REM duration profile. Example of early morning sleep onset resulting in a bimodal profile.

Within this structure, sleep cycles correspond to discrete energy levels: NREM duration and intensity map onto the period and amplitude of oscillatory movement between levels^1^. REM, by contrast, represents a delay in this propagation **–** a coherent superposition of sleep and wake states that allows for the controlled release of instability energy and the reduction of sleep need. REM intensity matches the energy released at each step to a lower level, while REM duration reflects the window of opportunity for that release **–** shaped by resonance. Importantly, the model curves for all four measures—NREM duration, NREM intensity, REM duration, and REM intensity—align closely with their empirical counterparts across sleep cycles, offering a precise mathematical fit to their distinctly different observed dynamics^1^.

We now show that the resonance that shapes REM episode duration is not singular but dual: it arises from two independently driven, overlapping forces **–** homeostatic and circadian (Fig. 1e,f). When aligned, they produce a typical bell-shaped REM curve; when misaligned, they give rise to linear or bimodal profiles. This dual-resonance structure explains longstanding empirical patterns under different conditions, including sleep deficit and sleep abundance. Importantly, this framework enables a physiologically grounded estimation of sleep deficit severity: not as hours missed, but as additional time required to return the system to equilibrium, derived from a single measurable parameter **–** REM episode duration.

This unified model not only reframes REM as a resonance-driven phenomenon, but also clarifies how homeostatic and circadian systems co-regulate the temporal structure of sleep. Together, these findings provide a new foundation for understanding REM dynamics across clinical, experimental, and ecological settings.

## RESULTS

### A dual-resonance mechanism shapes REM episode duration

Building on the validated core prediction of the wave model^1^ **–** that the characteristic bell-shaped trajectory of REM episode durations emerges from a resonance phenomenon near the equilibrium point between the sleep and wake potentials **–** we hypothesized that REM dynamics are shaped by two distinct but converging resonance processes: one homeostatic, the other circadian (Fig. 1e,f). In this dual-resonance framework, the homeostatic resonance curve is derived directly from the model’s internal dynamics and reflects the broader, composite nature of the homeostatic system—likely the result of multiple interacting physiological contributors. In contrast, the circadian resonance was modeled as a sharper, classical Lorentzian function, consistent with a single autonomous oscillator external to the energy landscape defined by the Morse potential. Fitting was performed by independently adjusting the amplitude and peak position of the homeostatic and circadian resonance curves. This allowed the data to determine the structure of the composite REM profile without bias toward expected timing or shape of the two resonance components.

To test whether this framework could explain REM architecture during regular sleep, we analyzed data from healthy young adults with no prior sleep debt and sleeping at their habitual times. The REM episode duration curve followed a smooth, near-symmetrical bell shape across cycles. Fitting the model to these data using both resonance components revealed a homeostatic peak earlier in the night and a circadian peak approximately 1.5 hours before awakening. Their relative time difference (Ψ = 2.9 h) and amplitudes produced a composite curve that matched the experimental REM durations with high fidelity (Fig. 2a).

**Figure 2.**
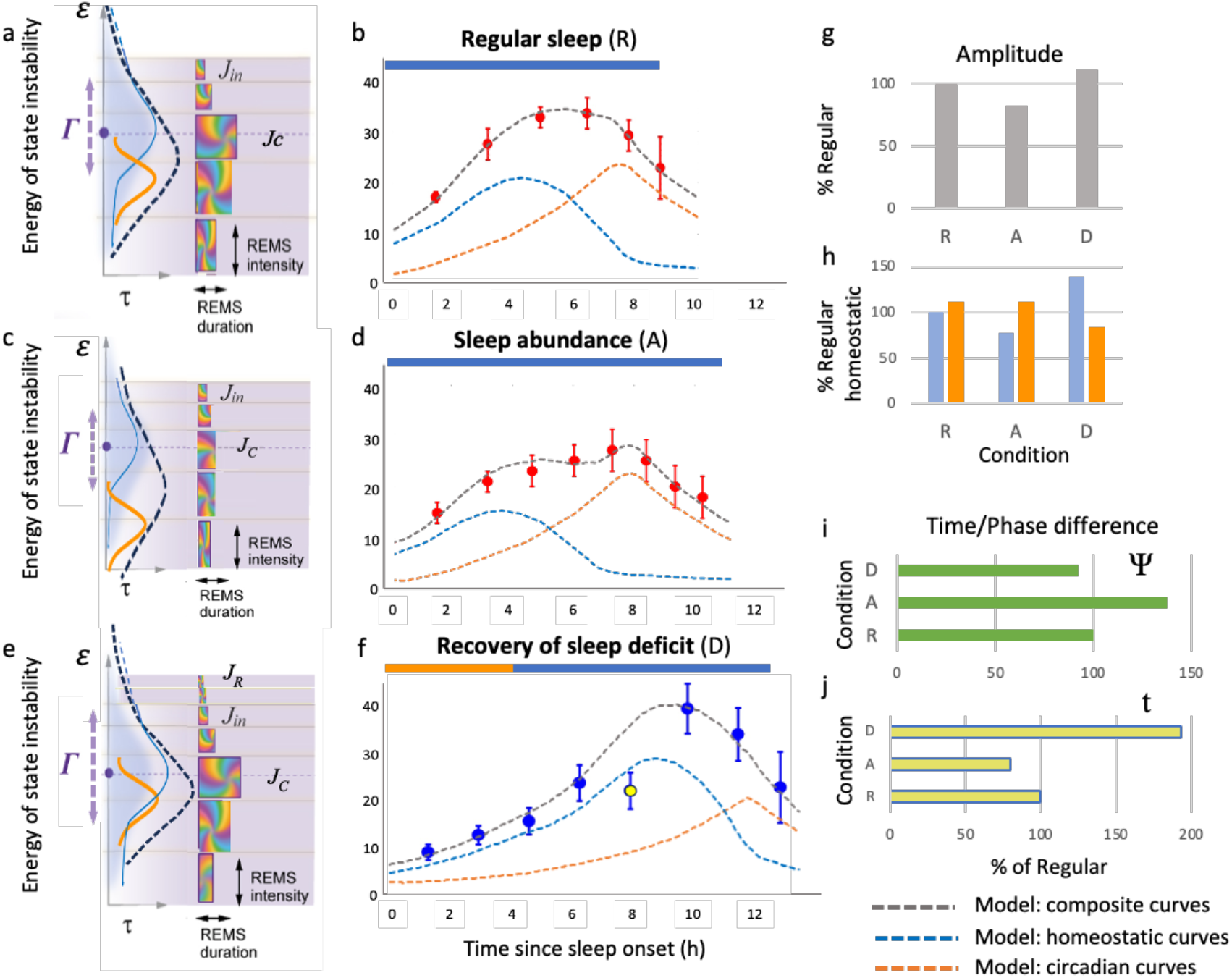
REM episode duration is shaped by homeostatic and circadian resonance, and modulated by sleep timing and deficit. (a, b) In baseline sleep, the model predicts a bell-shaped REM duration curve arising from synergistic alignment of homeostatic (blue) and circadian (orange) resonance. Experimental data match this profile (n = 39 nights, 24 subjects, mean ± SEM). (c, d) During chronic sleep abundance (14 h/night, 18:00–08:00 h, for 4 weeks; mean ± SEM), the homeostatic resonance peak is advanced and diminished, yielding a flattened REM profile (n = 268 nights, 11 subjects; mean ± SEM, see^2^). (e, f) Following 36 h of wakefulness, early sleep onset (19:00h) restores partial alignment between delayed homeostatic and circadian resonance, preserving the bell shape but with asymmetry (mean ± SEM, n = 9 nights, 9 subjects, see^6^). J_R_ - new initial level of recovery night. (g) Amplitude of composite REM profiles across regular (R, 100%), abundance (A), and deprivation (D) conditions. (h) Amplitude of homeostatic and circadian components, R-homeostatic =100%. (i) Phase difference (Ψ) between homeostatic and circadian peaks, R=100%. (j) Latency (t) from sleep onset to homeostatic resonance peak, R=100%. Horizontal bars in panels b,d,f show sleep periods (nighttime: blue; daytime: orange).

### Modulation of REM duration by sleep abundance and deficit

We next evaluated the model’s predictions on modified sleep schedules under entrained conditions that preserve the phase of the circadian resonance curve (Fig. 3a–g). Compared to regular sleep (Fig. 3a), delayed sleep onset increases state instability and reduces Ψ, thus increasing the overlap between homeostatic and circadian resonance curves (Fig. 3c). In contrast, earlier sleep onset – reflecting shorter prior wakefulness – lowers instability and increases Ψ (Fig. 3d). After prolonged wakefulness, sleep begins from a higher instability level (Fig. 3a), and changes in Ψ depend on timing of sleep initiation. Earlier sleep onset may allow for additional cycles above homeostatic equilibrium to occur prior to habitual bedtime, thus preserving the timing of the homeostatic resonance peak and Ψ (Fig. 3e), while regular bedtime reduces Ψ (Fig. 3f). A later bedtime can invert the alignment of the resonance curves, altering or preserving Ψ depending on their relative positions (Fig. 3g). These variations in Ψ affect the additive interaction of the resonance curves, modulating the amplitude of the composite signal and REM episode duration (Fig. 3b–g).

**Figure 3.**
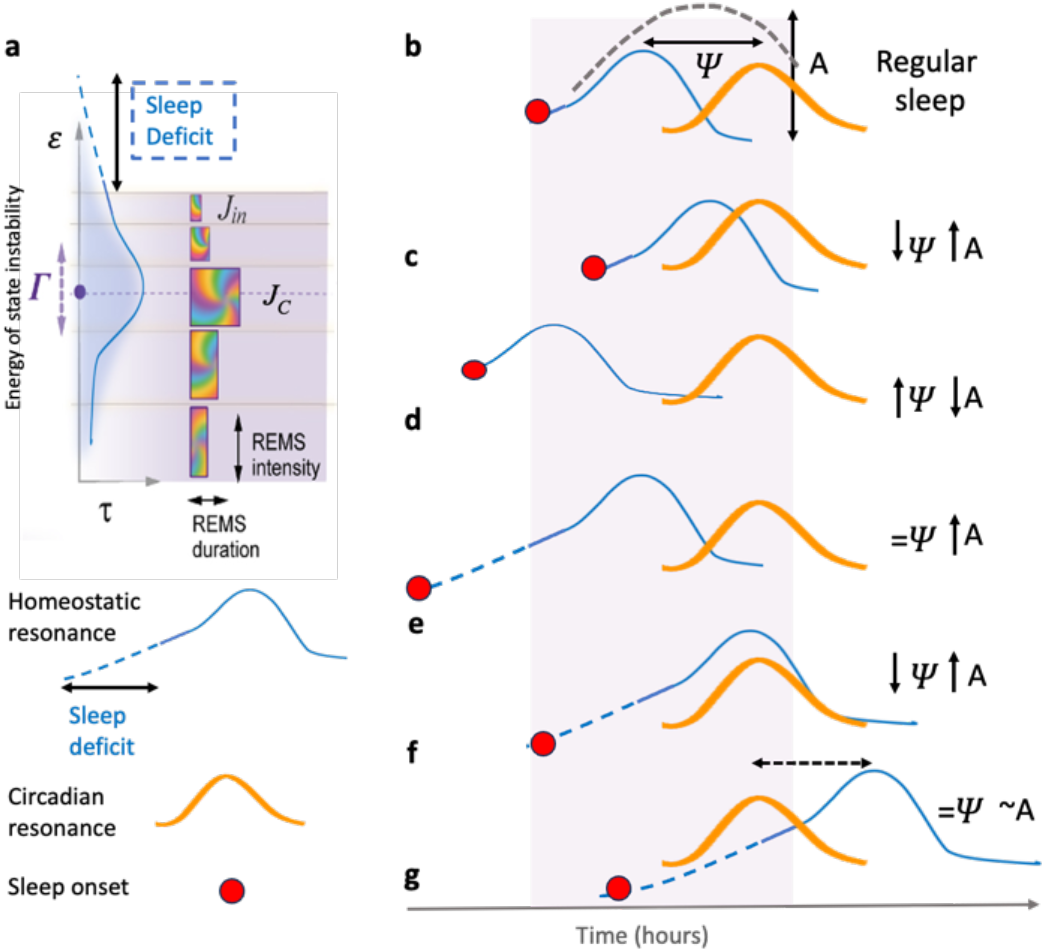
REM episode duration is shaped by the alignment between homeostatic and circadian resonance, and altered by sleep deficit. **(a)** The homeostatic resonance curve (solid blue) governs the timing of REM episodes via an energy-dependent resonance mechanism. With sleep deficit (dashed blue), the curve shifts upward to a higher energy level. **(b–g)** Schematic representations of different alignments between circadian (orange) and homeostatic (blue) resonance curves, influenced by sleep onset timing (red circles) and accumulated sleep deficit (dashed part of blue line). Small vertical arrows indicate changes in Ψ (the phase difference between resonance peaks) and in A, the amplitude of the composite resonance curve (black dashed line in **b**.) formed by the additive interaction of homeostatic and circadian components. Purple shading - regular sleep period. Under entrained conditions, the circadian resonance curve maintains a stable phase, as shown. Dashed arrow in **g**. – reversed order of homeostatic and circadian resonance.

To test these predictions, we analyzed REM episode durations from a chronic sleep extension protocol (14 h time-in-bed for four weeks)^2^. Previous modeling in this group identified a 13% widening of the Morse potential, consistent with a more spacious energy landscape, supporting more sleep cycles, smaller energy gaps, and reduced interaction strength between states^1^. As predicted by the lower energy of state instability accumulated over the 10-hour active period, the experimental REM profile was flatter and of lower amplitude, reflecting a reduced amplitude and extent of the homeostatic resonance curve (Fig. 2b, g, h). Despite the advance of the circadian resonance peak, which shifted further away from the final awakening, earlier bedtime and shorter latency from sleep onset to the homeostatic resonance peak (t, Fig. 2j) increased the phase difference (Ψ = 4.1 h, Fig. 2i). According to the model, increasing Ψ beyond 5 hours would be expected to produce a bimodal REM duration profile, as previously documented in forced desynchrony studies^3^.

To further test the model predictions, we then examined series of experiments aimed at extending sleep following sleep restriction or total deprivation with recovery sleep initiated at different times^4-6^. Consistent with the model’s prediction, sleep initiated in the morning exhibited a bimodal REM pattern4, with the model-estimated Ψ increasing to 6.8 h (Supplementary Fig. 1a,b). In contrast, when participants were given a 15-hour sleep opportunity initiated at midnight^5^, the REM duration curve showed a sharp, high-amplitude peak between 6 and 9 hours after sleep onset. The model reproduced this pattern by simulating a ∼2.5 h sleep deficit (Δt), which extended the homeostatic resonance curve and brought its peak into closer alignment with the circadian peak (i.e., reducing Ψ). This overlap amplified REM episode durations at mid-to-late night (Supplementary Fig. 1c-f), as predicted by the model (Fig. 3f).

Finally, we tested whether earlier sleep onset could buffer the effects of accumulated sleep deficit (Fig. 3e). In this condition, participants slept 4 hours earlier than usual following 36 hours of total wakefulness^6^. Despite the elevated sleep need, the REM duration profile retained its bell-shaped structure, albeit asymmetric, as expected. The model explained this by showing that the earlier sleep onset advanced the trajectory of the wavepacket (Fig. 3e), effectively compensating for the accumulated deficit (Δt ≈ 4.2 h). As a result, the homeostatic and circadian peaks remained properly aligned (Ψ ≈ baseline, Fig. 2i), preserving REM architecture despite substantial prior wakefulness. In these fits, the model also showed that homeostatic resonance amplitude increased with deficit, as was predicted by the model based on the energy-dependence of the strength of interaction between sleep and wake states (Fig. 2g, h). This is while the circadian component had reduced amplitude (Fig. 2h).

### Experimental REM profiles reveal a conserved resonance structure

We next asked whether acute sleep deprivation alters the homeostatic resonance structure itself or simply shifts the point at which the circadian system interacts with it. To test this, we compared REM episode durations during baseline and recovery sleep (Fig. 4a), and then normalized both to their respective maxima to control for amplitude differences (Fig. 4b). Strikingly, both profiles aligned with a single model-derived resonance curve (R^2^ > 0.98; Fig. 4c), indicating that the underlying structure remains stable across conditions.

**Figure 4.**
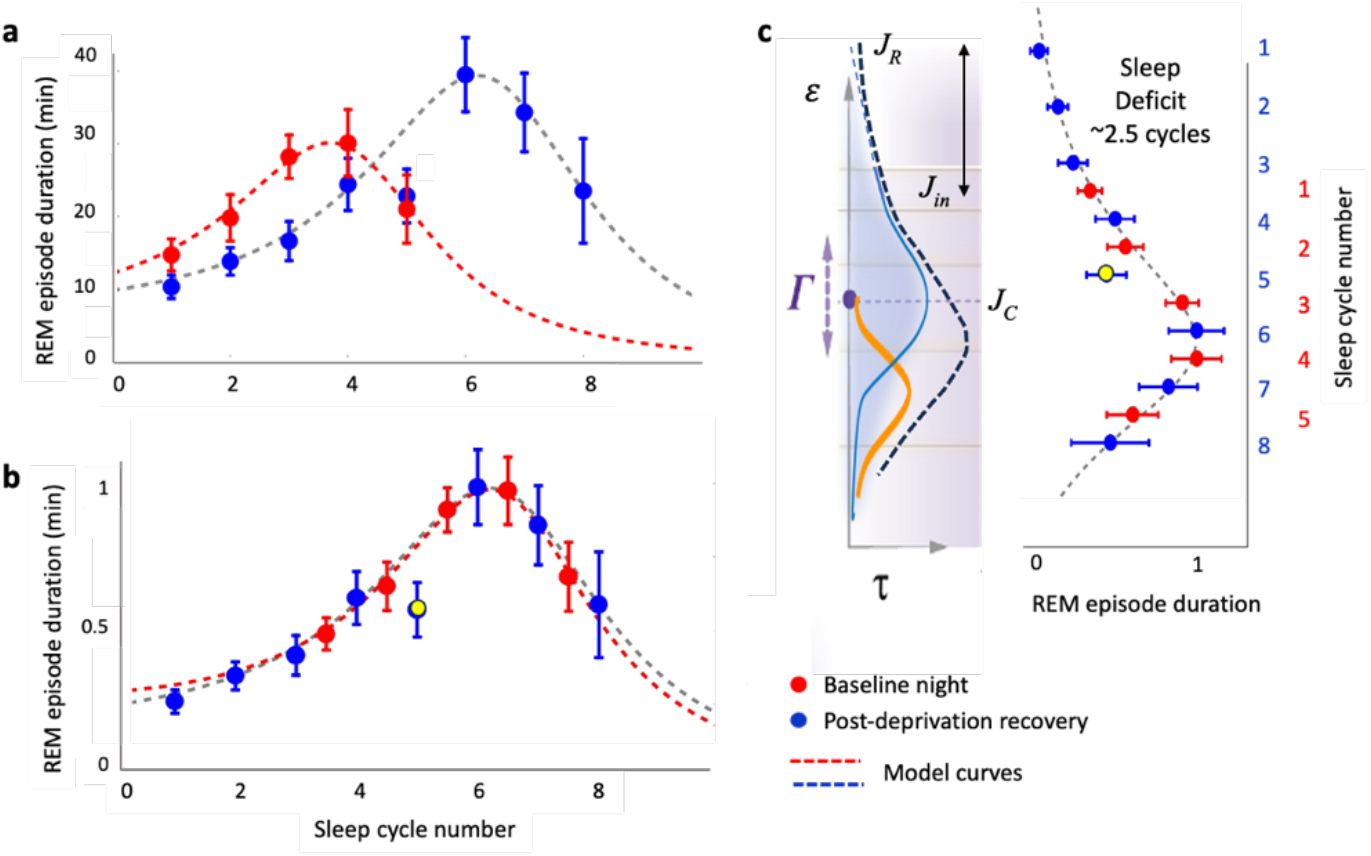
A conserved empirical resonance curve governs REM episode duration. (a) REM durations (mean ± SEM) during baseline and post-deprivation recovery sleep (n = 9; see^6^) plotted across successive cycles. Data closely match model-derived resonance curves. Cycle 5 of recovery sleep was excluded due to an adverse event affecting both NREM and REM duration. (b) REM durations normalized to individual group maxima reveal near-identical curve shapes across conditions, controlling for amplitude differences. (c) Left: Schematic illustrating the relative positions of homeostatic, circadian, and composite resonance curves in the presence of sleep deficit (dashed extension of blue line). The upward shift of ∼2.5 cycles between initial levels for recovery (*J*_*R*_) and for baseline (*Jin*) (black double-arrow) reflects the magnitude of additional state instability corresponding to sleep deficit. Right: The single blue model curve corresponds to recovery sleep; data as in **b**. This finding supports the existence of a stable, shared physiological structure across baseline and recovery sleep.

Crucially, the recovery curve exhibited an upward shift of approximately 2.5 cycles relative to baseline (Fig. 4c), validating the model-predicted sleep deficit of Δt ≈ 4.2 h. This estimate was based on the difference in the length of the homeostatic resonance curve required to reach equilibrium under regular (Fig. 2b) versus recovery conditions (Fig. 2f). This alignment provides empirical confirmation of a conserved energy architecture: a resonance structure that endures acute perturbation while precisely reflecting the system’s displaced entry point at recovery (*J*_*R*_).

### Phase-dependent REM duration patterns are altered by sleep deficit

Finally, we examined how REM episode duration patterns are modified when sleep is initiated at unusual circadian phases, a common misalignment challenge in shift work and jet lag^10^. Model simulations predicted that under entrained conditions without sleep deficit, REM profiles remain largely bell-shaped or, as in early morning sleep, bimodal, with timing and amplitude shifting across circadian phases (Fig. 5a). In contrast, when sleep begins with 2–4 h of accumulated deficit, thus delaying the homeostatic resonance peak (Fig. 3f,g), REM curves flatten and become progressively more linear within a standard 9-h sleep window (Fig. 5b,c). Extending sleep to 15 h (Fig. 5d) allows the system to reach the homeostatic equilibrium point, restoring the bell-shaped REM structure.

**Figure 5.**
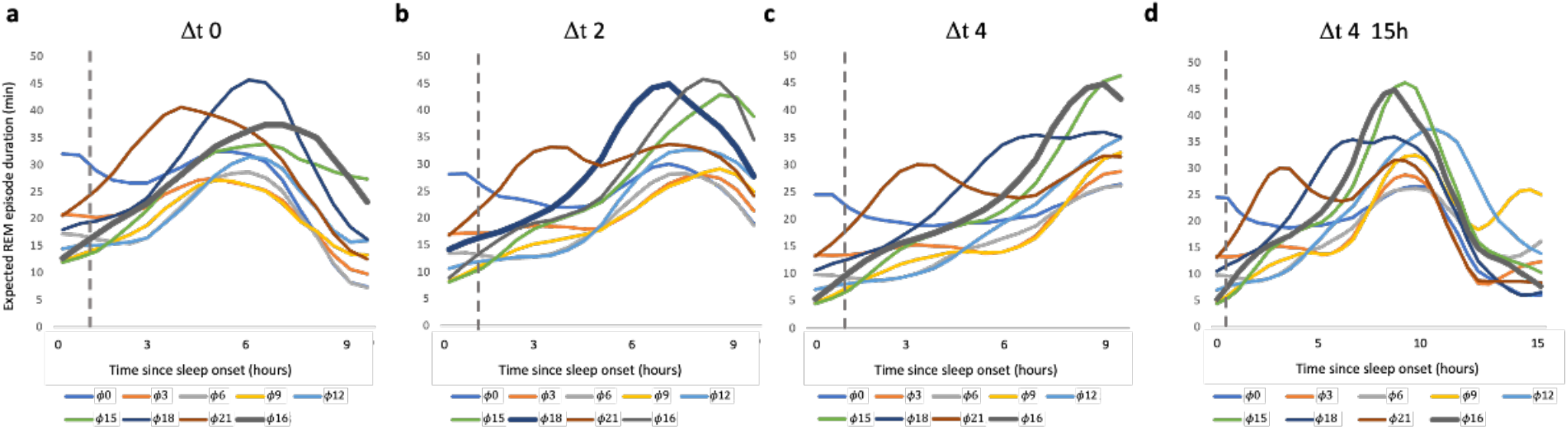
Circadian phase and sleep deficit jointly determine REM duration structure. (a) Model-predicted REM profiles across circadian phases with no sleep deficit. (b) With 2 h of deficit, curves shift and flatten. (c) With 4 h of deficit, most profiles become linear. (d) A 15 h sleep opportunity reveals bell-shaped REM profiles even with large deficit.

Φ-circadian phase of sleep onset relative to lights-on time (Φ0); Φ16 – in regular 16:8-h cycle (thick black line). Simulations illustrate how sleep deficit shifts homeostatic resonance timing beyond standard sleep windows.

The choice between absolute and relative values, as well as the length of the measurement frame, critically shapes the appearance of REM duration curves. The model profiles reflect REM episode durations measured in absolute terms, which show some differences from results reported in forced desynchrony protocols, where REM is often expressed relative to total sleep time or recording duration^3^. We use absolute values because concurrent changes in NREM duration and intermittent awakenings can obscure the underlying structure of REM when expressed as proportions (Supplementary Fig. 2a,b). This may also explain the earlier peak of the circadian resonance curve – around the core body temperature minimum – compared to the end-of-sleep REM peak typically reported in forced desynchrony studies^3^ (Supplementary Fig. 2c). Moreover, absolute REM duration is critical for identifying the timing and magnitude of homeostatic resonance expression—an effect essential for estimating sleep deficit.

Together, these results show that REM episode duration reflects a dynamic interaction between two independent resonance processes. The amplitude and phase alignment of homeostatic and circadian systems determine how REM is expressed across diverse physiological and environmental contexts.

## DISCUSSION

In the wave model framework, physiological instability accumulates during wakefulness and is progressively resolved through structured NREM reorganization and released during REM. This evolution, modeled as probability waves propagating within a Morse potential, generates cyclic NREM–REM architecture through interaction between sleep and wake states. Critically, REM duration reflects not only interaction strength, but also resonance between states, which prolongs overlap, stabilizes superposition, and provides time for efficient instability release at the end of each sleep cycle (Fig. 1c).

In addition to the precise mathematical description of NREM and REM dynamics previously demonstrated^1^, this study offers strong empirical confirmation of the resonance hypothesis. The data reveal a single, conserved resonance curve—maintained across both baseline and recovery sleep (Fig. 4). While sleep deprivation shifts the wavepacket’s entry point by raising state instability at sleep onset, the shape and position of the resonance remain unchanged **–** a hallmark of robust physiological architecture. Furthermore, comparison of baseline and recovery curves quantifies the additional sleep cycles required to compensate for sleep deficit accrued during extended wakefulness.

This work further advances the wave model by demonstrating that resonance is not singular but dual: shaped by independent yet converging homeostatic and circadian components. While both systems have long been implicated in REM regulation^3^, our findings show how their interaction determines REM duration patterns across sleep. The relative phase and amplitude of these two components shape the composite resonance curve and REM duration profile, producing bell-shaped, linear, or bimodal forms depending on sleep timing and accumulated deficit.

The homeostatic resonance reflects a state-dependent capacity to sustain REM, peaking near the system’s energy equilibrium (Fig. 1c). As a result, it does more than define REM structure **–** it provides a direct, quantitative marker of sleep deficit in physiologically meaningful units: not merely hours lost, but the additional hours or cycles required to return to homeostasis. Notably, this estimate does not require baseline comparison, as the shape of the homeostatic resonance curve itself reveals the remaining distance to equilibrium. Together, REM duration offers a dynamic readout of physiological strain and recovery progress, reinforcing emerging proposals that REM may serve as a sensitive index of sleep quality^11^.

In parallel, the position of the circadian resonance peak **–** also revealed by the REM duration curve **–** offers complementary insight. First, it reflects the timing of circadian phase relative to clock time, providing an accessible measure of phase position and potential misalignment with environmental zeitgebers. Second, the phase relationship between homeostatic and circadian peaks defines the degree of intrinsic misalignment between the sleep state and circadian system itself. In both cases, these insights may guide targeted strategies to restore circadian harmony in conditions such as jet lag, shift work, or circadian rhythm disorders.

During regular sleep, circadian influence plays a key role in amplifying and extending REM toward the end of the night **–** a mechanism that enhances instability clearance as homeostatic drive wanes. In our model, REM intensity reflects the energy released during level transitions, which rises steadily toward morning, while REM duration reflects the available release window, governed by resonance alignment. When circadian and homeostatic peaks coincide late in sleep, REM expression is maximized to support full recovery.

This synergy enhances sleep–wake resilience. By enabling deeper instability clearance at night and stabilizing daytime wakefulness, the circadian system broadens the homeostatic system’s dynamic range. The organism enters wakefulness from a lower-instability baseline and can remain further from collapse, tolerating greater wake-state instability before requiring sleep onset (Fig. 1a,b). These complementary dynamics extend both the amplitude and flexibility of the sleep–wake cycle.

Taken together, the model offers a mechanistic explanation for how homeostatic and circadian systems jointly regulate REM. By identifying their distinct yet interacting roles **–** homeostatic resonance as dynamic instability management, circadian resonance as temporal modulation **–** we provide a unified framework for understanding sleep architecture as an adaptive, resonance-driven process. This perspective not only reframes REM as a physical phenomenon, but also introduces new opportunities to use REM structure as a sensitive tool for sleep medicine, shift-work adaptation, and chronobiological research.

## METHODS

### Sleep Data Collection

Modeling of REM episode duration was based on data collected from healthy young adults, as previously described^1^. The dataset was part of a broader investigation into the circadian regulation of sleep and hormonal rhythms (“Multimodal Circadian Rhythm Evaluation”; PI: IVZ, funded by Pfizer Inc.). The study was conducted in accordance with the Declaration of Helsinki and approved by the Boston University Institutional Review Board. All participants provided written informed consent.

Twenty-four healthy male volunteers (mean ± SEM: 24.5 ± 4.4 years; range: 19–34) were included. Participants reported habitual sleep durations of 7–9 hours per night, with <1.5 h variability on weekends, no history of sleep disorders or chronic health conditions, no regular medications, and no recent trans-meridian travel. All were non-smokers and consumed <3 cups of coffee per day.

The study consisted of three phases:

- **Phase 1:** Participants selected and maintained a stable sleep/wake schedule for five days, verified via actigraphy and wearable devices. Those demonstrating high compliance proceeded to Phase 2.
- **Phase 2:** Two weeks of outpatient monitoring using multiple actigraphy devices and an Oura ring for continuous recordings of activity, skin temperature, and heart rate.
- **Phase 3:** Three nights of inpatient polysomnography (PSG), core body temperature recordings, and saliva/blood sampling (night 3) under controlled conditions.

Only nights 1 and 2 with sleep efficiency ≥85% and no evidence of sleep pathology were included (n = 39). Sleep episodes were visually scored in 30-second epochs. NREM–REM cycles were defined as a sequence of NREM sleep (≥10 minutes) followed by REM sleep (≥3 minutes). Intermittent wakefulness was excluded from cycle durations and analyzed separately.

### Mathematical Modeling of REM Duration

#### Wave Model Framework and Homeostatic Resonance

This work builds upon a previously introduced wave model of sleep dynamics^1^, in which sleep and wake states are represented as interacting probability waves propagating through separate interacting Morse potentials.

- **NREM episodes** correspond to wavepacket oscillations within the sleep potential well. Their duration is proportional to the period of oscillation, and their intensity (slow-wave activity) scales with the square of the oscillation amplitude.
- **REM episodes** correspond to coherent superposition of wake and sleep waves during the transitions between adjacent energy levels, with REM intensity proportional to the energy gap (Δε), and duration determined by resonance strength, with resonance level (*Jc*) positioned near the homeostatic equilibrium region.

NREM durations for the n-th cycle are modeled as:

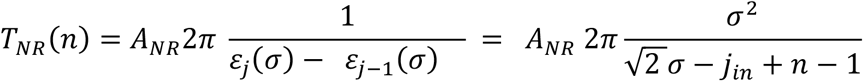

and REM durations as:

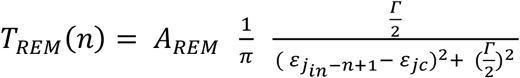

where σ is the characteristic width of the Morse potential, *Γ* is the resonance width, and *A*_*NR*_ and *A*_*REM*_ are scaling constants to convert relative model values to time in minutes.

### Dual-Resonance Model for REM Duration

To account for circadian influences, REM durations were further modeled using a dual-resonance framework:

- **Homeostatic resonance** retained the Gaussian-based structure derived from the wave model.
- **Circadian resonance** was modeled as an independent Lorentzian curve, reflecting a stable oscillator phase-locked to the circadian cycle.

REM duration at each cycle was modeled as:

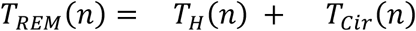

With:

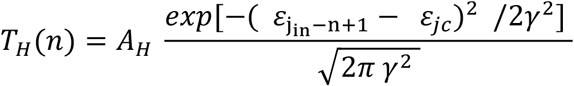

And

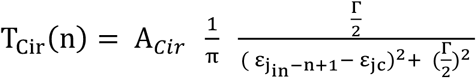

Here, where *Γ* and *γ* are the resonance widths for homeostatic and circadian components, and *A*_*H*_ and *A*_*Cir*_ are normalization constants.

### Model Fitting Strategy

Model fitting involved independent adjustment of the amplitude and timing of each resonance curve. The homeostatic curve structure was preserved from prior work^1^, with no optimization of *Jin* or resonance width. This approach ensured internal consistency across datasets (regular sleep, extended sleep, and recovery following sleep deprivation) and avoided bias toward expected peak timing.

## Data availability statement

The dataset generated by the co-author (IVZ) and analyzed during the current study is available from the corresponding authors on reasonable request.

The mathematical algorithms of the wave model of sleep dynamics in “Wolfram Mathematica” format are available from the corresponding author (VK) on reasonable request.

## Author contributions

V.K., and I.V.Z. designed research; I.V.Z performed sleep research; V.K. conducted mathematical modeling; V.K. and I.V.Z. wrote the paper.

## Supplementary Materials

**1. Figure S1** Circadian–homeostatic alignment shapes REM profiles under altered sleep timing and history.

**2. Figure S2** Absolute and relative REM measures result in different patterns.

**Supplementary Figure 1.**
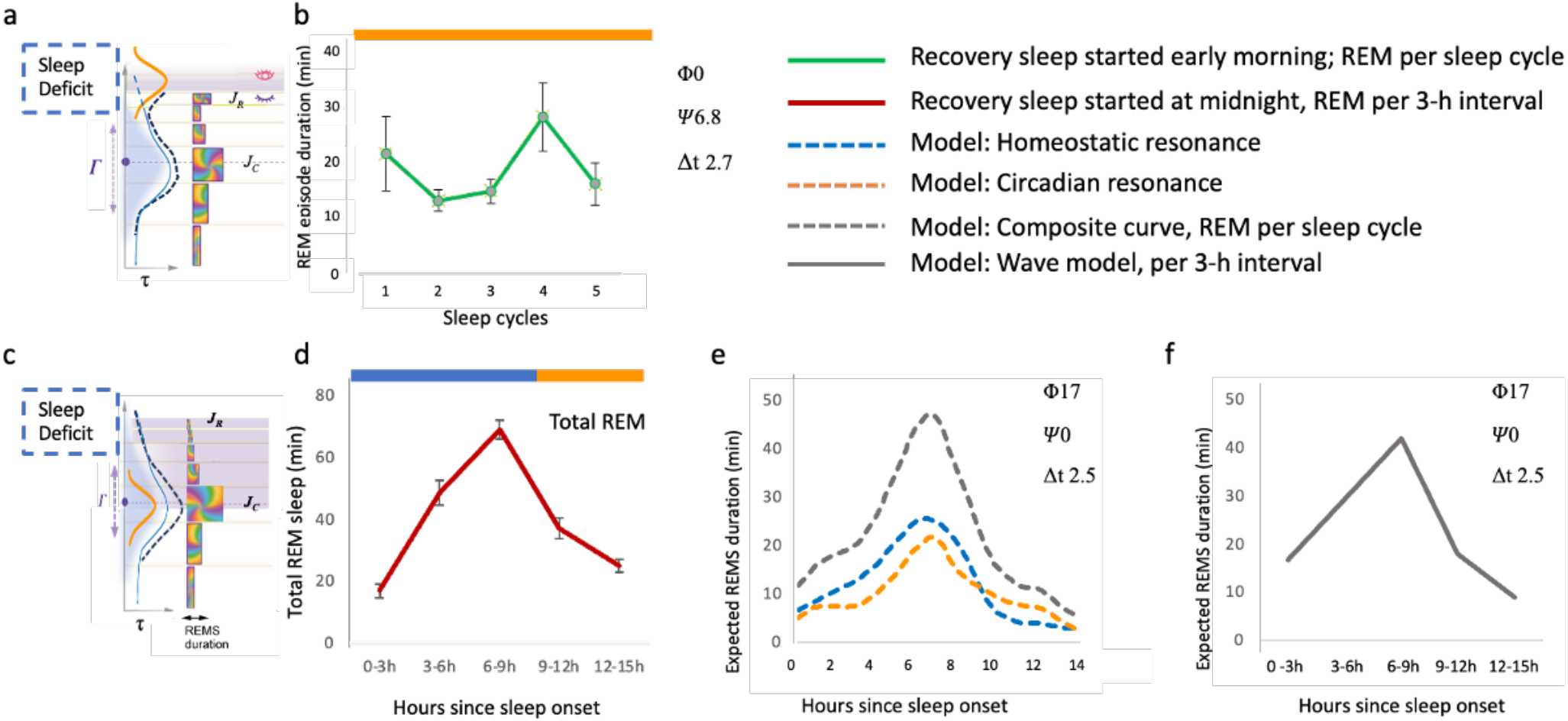
Circadian–homeostatic alignment shapes REM profiles under altered sleep timing and history. a. Schematics of dissociation between homeostatic and circadian resonance curves due to early morning sleep onset, with sleep deficit following overnight sleep deprivation, predicts a bimodal REM duration profile. b. Bimodal REM episode duration profile of recovery sleep initiated in the morning, at 07:00h, following 24-h wakefulness. Data: mean (SEM), *n* = 8, adapted from Table 1, Dijk et al., 1991^4^. High data variability and only 4-5 subjects completing cycles 3-5 precluded precise model analysis. Estimated phase difference between circadian and homeostatic resonance peaks –∼Ψ6.8h, sleep deficit ∼Δt2.7h. Φ0 ∼ habitual lights-on time. c. Schematics of reduced phase difference between homeostatic and circadian resonance due to post-deprivation shift in homeostatic resonance curve resulting in the overlap of two resonance peaks. This leads to higher and sharper than usual peak in REM duration. d. Total REM duration per 3-hour interval during a 15-hour recovery sleep opportunity initiated at midnight, following multiple days of sleep restriction. Data: mean (SEM), *n* = 9, adapted from Table 1, Dijk et al., 1991^5^. e. Model simulation of REM episode duration based on the 2.5-hour sleep deficit (Δt 2.5), Ψ0h, and sleep onset at circadian phase Φ17 (Φ0∼lights-on time). Curves: blue – homeostatic resonance, orange – circadian resonance, black – predicted composite REM duration. *Note:* This simulation is included for illustrative purposes, as the original data were aggregated into 3-hour intervals, precluding formal model fitting. f. Predicted REM episode durations from b., summed across 3-hour intervals to match panel a.

**Supplementary Figure 2.**
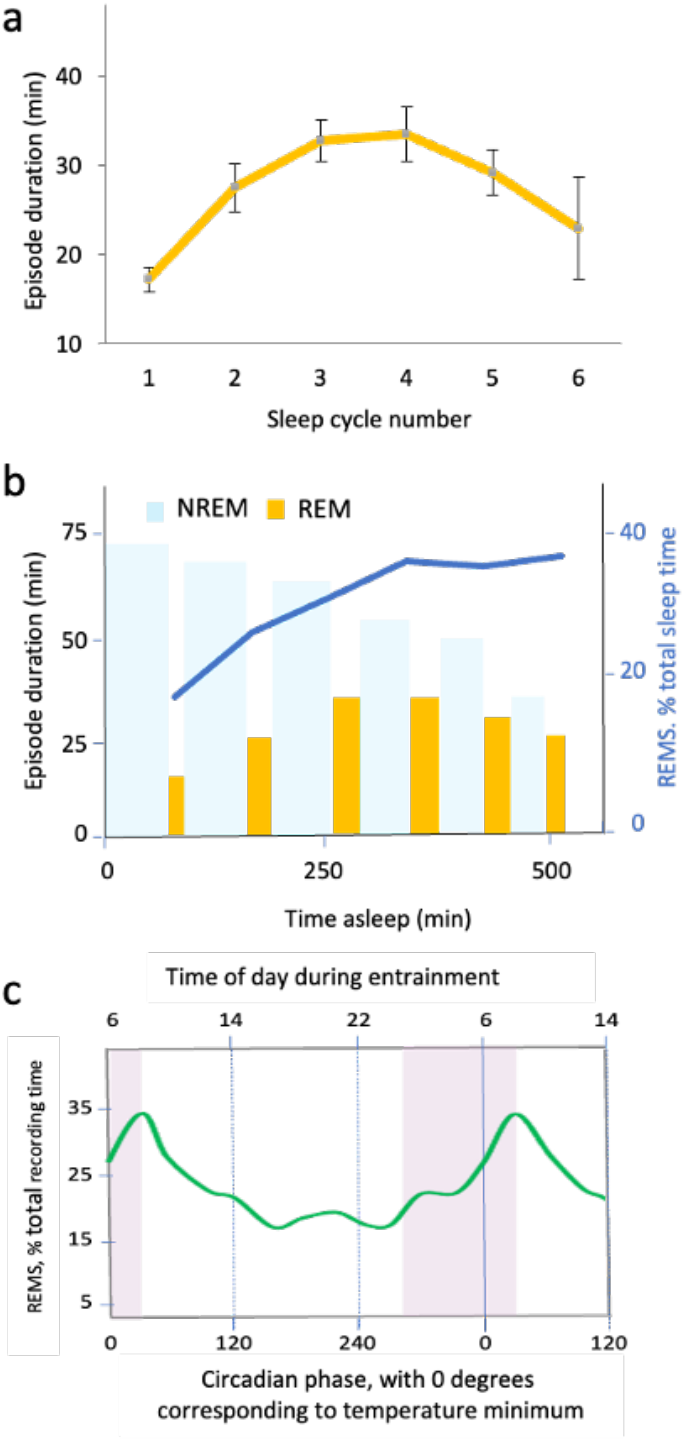
Absolute and relative REM measures result in different patterns. a. REM durations plotted by sleep cycle (absolute values) show a bell-shaped curve (n=39 nights). b. Relative REM (% of total sleep time, blue line) shows a monotonic increase due to declining NREM duration (blue bars) across cycles (n=39 nights). Yellow bars – REM episode duration, as in **a.** c. Circadian phase–dependent REM duration curve based on %REM per Recording time (RT) peaks near habitual wake time, ∼1–2 h after the core body temperature minimum (from forced desynchrony studies, Figure 3b in^3^). In contrast, the wave model predictions based on data expressed in absolute values of REM duration per sleep cycle place the peak of the circadian resonance curve that modulates REM episode duration near the temperature minimum, or Phase 0 on this plot (see Fig. 2 of Main text).

